# Coordination Among Multiple Receptor Tyrosine Kinase Signals Controls *Drosophila* Developmental Timing and Body Size

**DOI:** 10.1101/2020.09.01.278382

**Authors:** Xueyang Pan, Michael B. O’Connor

## Abstract

Body size and the timing of metamorphosis are two important interlinked life-history traits that define holometabolous insect development. Metamorphic timing is largely controlled by a neuroendocrine signaling axis composed of the prothoracic gland (PG) and its presynaptic neurons (PGNs). The PGNs produce prothoracicotropic hormone (PTTH) that stimulates the PG to produce the metamorphosis inducing hormone ecdysone (E) through activation of Torso a Receptor tyrosine kinase the Receptor Tyrosine kinase and its downstream Ras/Erk signal transducers. Here we identify two additional timing signals produced by the RTKs Anaplastic lymphoma kinase (Alk) and the PDGF/VEGF-receptor related (PvR), Similar to Torso, both Alk and PvR trigger Ras/Erk signaling in the PG to up regulate expression of E biosynthetic enzymes, while Alk also suppresses autophagy induction after critical weight by activating Pi3K/Akt. When overexpressed, both RTKs hyperactivate an endogenous low-level Jak/Stat signal in the PG resulting in developmental delay or arrest. The Alk ligand Jelly belly (Jeb) is produced by the PGNs, and together with PTTH serves as a second PGN derived tropic factor that stimulates E production by the PG. In addition, we find that Pvf3, a PvR ligand, is also produced by the PGNs, but we show that the activation of PvR primarily relies on autocrine signaling by PG-derived Pvf2 and Pvf3. These findings illustrate that a multitude of juxtracrine and autocrine signaling systems have evolved to regulate the timing of metamorphosis, the defining event of holometabolous development.

## Introduction

Body size is one of the most important traits of a multicellular organism. In species whose growth is determinate, the body growth of an individual is largely completed when it matures into an adult (Callier and Nijhout, 2013). A good example of determinate growth is found among holometabolous insects, such as the fruit fly *Drosophila melanogaster*. During development, the size of a *Drosophila* larva increases one hundred-fold during its three molts, but does not change after metamorphosis, the developmental stage that transitions the juvenile larval form into the sexually mature adult fly. Therefore, the control of metamorphic timing is a key factor that regulates final body size.

In the past decades, numerous studies in *Drosophila* and other holometabolous insect species have demonstrated that the onset of metamorphosis is regulated through a neuroendocrine signaling axis composed of two central information processing nodes: The prothoracic gland (PG) which produces the metamorphosis inducing steroid hormone ecdysone (E), and a bilateral pair of brain neurons (PGNs) that innervate the PG and release the neuropeptide PPTH that stimulates E production. (McBrayer et al., 2007; Yamanaka et al., 2015; Yamanaka et al., 2013a). After release into the hemolymph, E is taken up by peripheral larval tissues through a specific importer (ECI) and then converted into its active form, 20-hydroxyecdysone (20E) by the enzyme Shade (Okamoto et al., 2018; Petryk et al., 2003) Subsequently, 20E stimulates metamorphosis via activation of the ecdysone receptor complex (EcR)/ultraspiracle (Usp) and stimulation of tissue-specific downstream transcriptional cascades (Hill et al., 2013).

In this scheme, PTTH functions as a trophic hormone to stimulate PG growth and E synthesis (Shimell et al., 2018; Smith and Rybczynski, 2012). In PG cells, PTTH binds to Torso, a receptor tyrosine kinase (RTK) family member, and stimulates the E biosynthetic pathway via Ras/Erk signaling (Rewitz et al., 2009). As the two central nodes on the neuroendocrine axis, both the PG and the PGNs receive additional diverse internal and external signals modulate their output appropriately. For instance, the PG cells respond to insulin signals reflecting the general nutritional state (Colombani et al., 2005; Mirth et al., 2005). In addition, systemic BMP signals help coordinate metamorphosis with appropriate imaginal disc growth (Setiawan et al., 2018). The PGNs in turn, receive presynaptic inputs from various upstream neurons that regulate circadian and pupation behaviors (Deveci et al., 2019; Imura et al., 2020; McBrayer et al., 2007). They also respond to tissue damage signals to delay maturation onset until the damage is resolved (Colombani et al., 2015; Colombani et al., 2012; Garelli et al., 2012; Garelli et al., 2015; Vallejo et al., 2015).

Although it is widely accepted that PTTH is the key neuropeptide that triggers developmental maturation in holometabolous insects (Deveci et al., 2019; McBrayer et al., 2007; Shimell et al., 2018; Smith and Rybczynski, 2012), several studies indicate that additional timing signals are also likely. The first suggestion that PTTH is not the sole timing signal came from PGN ablation studies in *Drosophila* where it was found that up to 50% of animals with no PGNs still undergo metamorphosis, but after a prolonged ∼5 day developmental delay (Ghosh et al., 2010; McBrayer et al., 2007). Subsequently, it was found that genetic null mutations in *Drosophila* PTTH gene only produced a one-day developmental delay and had little effect on viability (Shimell et al., 2018). In this case electrical stimulation of the mutant PGNs restore proper timing while inactivation produced a more substantial 2-day delay (Shimell et al., 2018). Ptth null mutants have also been generated in *Bombyx mori* and while most animals do arrest development a fraction still escape and produce adults (Uchibori-Asano et al., 2017). Taken together, these studies strongly indicated that the PGNs produce additional timing signals besides PTTH.

RTK family receptors has been speculated to mediate the additional PGN signal, since blocking Ras/Erk pathway in the PG causes strong developmental defects, phenocopying the PGN ablation model rather than the *ptth* mutant (Cruz et al., 2020; Rewitz et al., 2009). Epidermal growth factor receptor (Egfr) has recently been demonstrated crucial for PG tissue growth, ecdysone synthesis and secretion. However, the Egfr pathway is activated by autocrinal signals from the PG, which does not involve the activity of PGNs (Cruz et al., 2020). In the present study, we identify two additional RTK family receptors, Anaplastic lymphoma kinase (Alk) and PDGF- and VEGF-receptor related (Pvr), which play important roles in the PG controlling metamorphic timing. Interestingly, the Alk ligand Jelly belly (Jeb) and Pvr ligand Pvf3 are expressed in the PGNs, verifying that the prothoracicotropic function of PGNs is mediated by multiple signaling molecules.

## Results

### Targeted screening of *Drosophila* RTKs for factors controlling developmental timing

Based on the speculation that RTKs could mediate the trophic signals from PGNs to the PG, we performed a targeted RNAi screening using PG-specific *phm-Gal4* driver to identify RTKs in the PG that regulate the timing of pupariation. Since the knockdown efficiency of RNAi construct varies, we carried out the screening using RNAi lines from the Transgenic RNAi Project (TRiP) and compared the results with recently published genome-wide screening using RNAi lines from Vienna Drosophila Resource Center (VDRC) (Danielsen et al., 2016). Insulin receptor (InR) and Torso, whose functions in the PG have already been readily documented (Colombani et al., 2005; Mirth et al., 2005; Rewitz et al., 2009), were identified in both screens. In addition, we found Alk and Egfr as hits in our TRiP screen while Pvr was a potential hit in the previous genome wide screen (Table S1). Since the role of Egfr in the PG has been documented in a recent study (Cruz et al., 2020), in this report we focused our efforts of elucidating the roles of Alk and Pvr in regulation of metamorphic timing and body size.

### Alk and Pvr are required for normal metamorphic timing and body size control

Following the initial screen, we first sought to verify the developmental timing phenotype of Alk and Pvr suppression larvae using multiple RNAi constructs as well as dominant negative receptors. In line with the screening result, knocking down Alk in the PG using two different RNAi constructs caused delay of developmental timing. Furthermore, overexpressing dominant negative Alk resulted in developmental arrest in the L3 stage (Figure 1A and S1A). Similarly, the developmental delay phenotype of Pvr suppression larvae was produced by two independent Pvr RNAi constructs and a third produced developmental arrest. In addition, expression of a dominant negative Pvr in the PG also produced developmental delay (Figure 1A and S1B).

**Figure 1.**
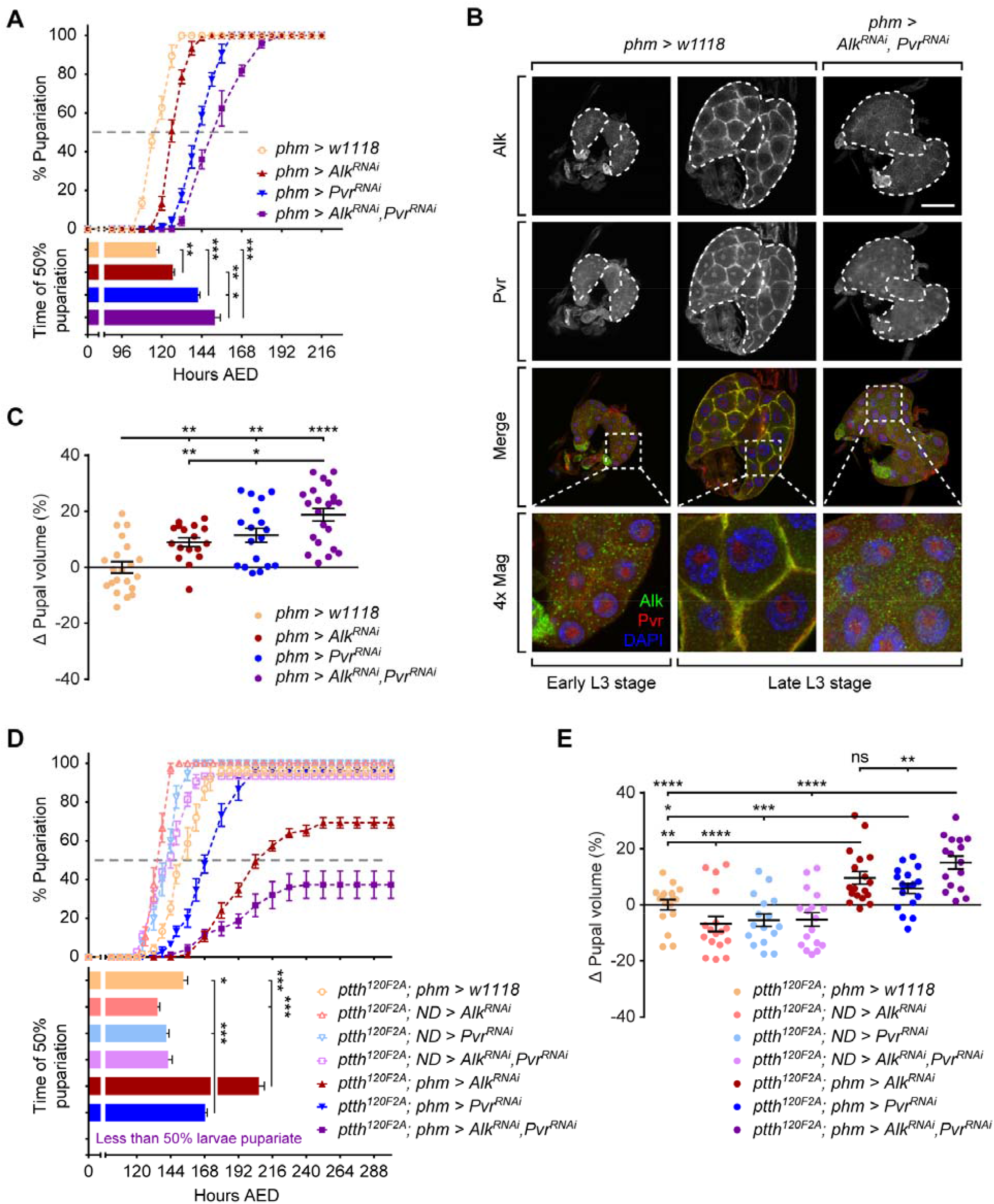
Alk and Pvr regulate developmental timing and body size in coordination with PTTH/Torso pathway. (A) Pupariation timing curves and the time of 50% pupariation of *phm*>*w1118, phm*>*Alk*^*RNAi*^, *phm*>*Pvr*^*RNAi*^ and *phm*>*Alk*^*RNAi*^, *Pvr*^*RNAi*^ larvae. (B) Immunofluorescence images of *phm*>*w1118* and *phm*>*Alk*^*RNAi*^, *Pvr*^*RNAi*^ PGs stained with anti-Alk and anti-Pvr antibodies. Dash lines outline the PG area of the ring gland. Scale bar, 50μm. (C) Relevant pupal volume changes in animals tested in (A). (D) Pupariation timing curves and the time of 50% pupariation of *phm*>*w1118, phm*>*Alk*^*RNAi*^, *phm*>*Pvr*^*RNAi*^ and *phm*>*Alk*^*RNAi*^, *Pvr*^*RNAi*^ larvae with *ptth*^*120F2A*^ mutant background. ND, no driver. (E) Relevant pupal volume changes in animals tested in (D). (A, C-E) Mean ± SEM; p values by unpaired t-test (n=3 in A and D, n=17-22 in C and E; ns, not significant, *p<0.05, **p<0.01, ***p<0.001, ****p<0.0001).

Using immunofluorescent staining, we examined the expression of Alk and Pvr in the PG and tested knockdown efficiency of the RNAi lines used above. In control larvae, strong expression of Alk and Pvr was observed in the PG of late-L3 stage larvae, reflected by the distinct fluorescence signals on PG cell membrane (Figure 1B). Interestingly, the expression of both receptors was remarkably weaker in early-L3 stage (Figure 1B), indicating that the signal outputs from these receptors may be stronger in late-L3 stage when larvae approach the onset of metamorphosis. When expressing RNAi lines (Alk RNAi #1 and Pvr RNAi #2) (Figure S1A-B), we found that the expression of both receptors in the PG was effectively depleted in late-L3 stage (Figure 1B). Since these RNAi constructs induce efficient gene knockdown, we used them in our following studies. When knocking down either Alk or Pvr alone, we observed minor developmental delay. However, simultaneously knockdown of both receptors by *phm*>*Alk*^*RNAi*^, *Pvr*^*RNAi*^ leads to a more prolonged developmental delay (Figure 1A). Thus, we conclude that both Alk and Pvr act in the PG to regulate metamorphic timing perhaps in an additive manner. As for the developmental arrest phenotype observed in other crosses (*phm*>*Alk*^*DN*^ and *phm*>*Pvr*^*RNAi #3*^) (Figure S1A-B), we speculate that they may result from unknown detrimental effects from the transgenes or the genetic background of these lines.

In addition to timing, we measured the pupal size of Alk and Pvr suppression animals. The size of *phm*>*Alk*^*RNAi*^ and *phm*>*Pvr*^*RNAi*^ pupae are larger than that of the *phm*>*w1118* controls, while the *phm*>*Alk*^*RNAi*^, *Pvr*^*RNAi*^ animals formed pupae of even larger size (Figure 1C). We conclude that both Alk and Pvr are required in the PG for normal developmental timing and body size control.

### Loss of Alk and Pvr causes stronger developmental defects in *ptth* mutants

The mild developmental delay phenotype of Alk and Pvr suppression animals is comparable to that of *ptth* mutants (Shimell et al., 2018). Since Alk, Pvr and Torso are all RTKs, we propose that the Alk and Pvr pathways may function additively or synergistically with PTTH/Torso pathway to control developmental timing. To test this possibility, we knocked down Alk and Pvr in the PG of *ptth* mutants and examined whether the timing of pupariation is further prolonged. Consistent with our conjecture, both *ptth; phm*>*Alk*^*RNAi*^ and *ptth; phm*>*Pvr*^*RNAi*^ larvae took longer to pupariate than the *phm-Gal4* and no driver (*ND*) controls and 30% of *ptth; phm*>*Alk*^*RNAi*^ larvae even failed to pupariate (Figure 1D). Moreover, longer developmental delay and higher rates of developmental arrest at the L3 stage were observed in *ptth; phm*>*Alk*^*RNAi*^, *Pvr*^*RNAi*^ larvae in which all three RTK pathways were suppressed (Figure 1D). In parallel to the developmental timing change, the pupal size of double or triple RTK suppression animals were also larger than controls (Figure 1E). These data demonstrate that the Alk and Pvr work in association with PTTH/Torso and suggest that the receptors may share the same downstream signaling pathway to regulate developmental timing.

### Alk and Pvr facilitate ecdysone synthesis and Halloween gene expression by activating Ras/Erk pathway

It is well established that PTTH/Torso signaling facilitates pupariation activity by stimulating ecdysone synthesis in the PG via Ras/Erk pathway (Rewitz et al., 2009). To determine whether Alk and Pvr function via the same mechanism, we first examined the ecdysone level in Alk and Pvr suppression larvae. In mid-L3 stage, we did not observe a significant difference in the ecdysone level among *phm*>*w1118, phm*>*Alk*^*RNAi*^, *phm*>*Pvr*^*RNAi*^ and *phm*>*Alk*^*RNAi*^, *Pvr*^*RNAi*^ animals. However, at the time point when *phm*>*w1118* larvae are at the wandering stage the receptor suppression larvae produce a markedly lower level of ecdysone than *phm*>*w1118* controls (Figure 2A), suggesting that the ecdysone synthesis is compromised when Alk and/or Pvr is suppressed in the PG.

**Figure 2.**
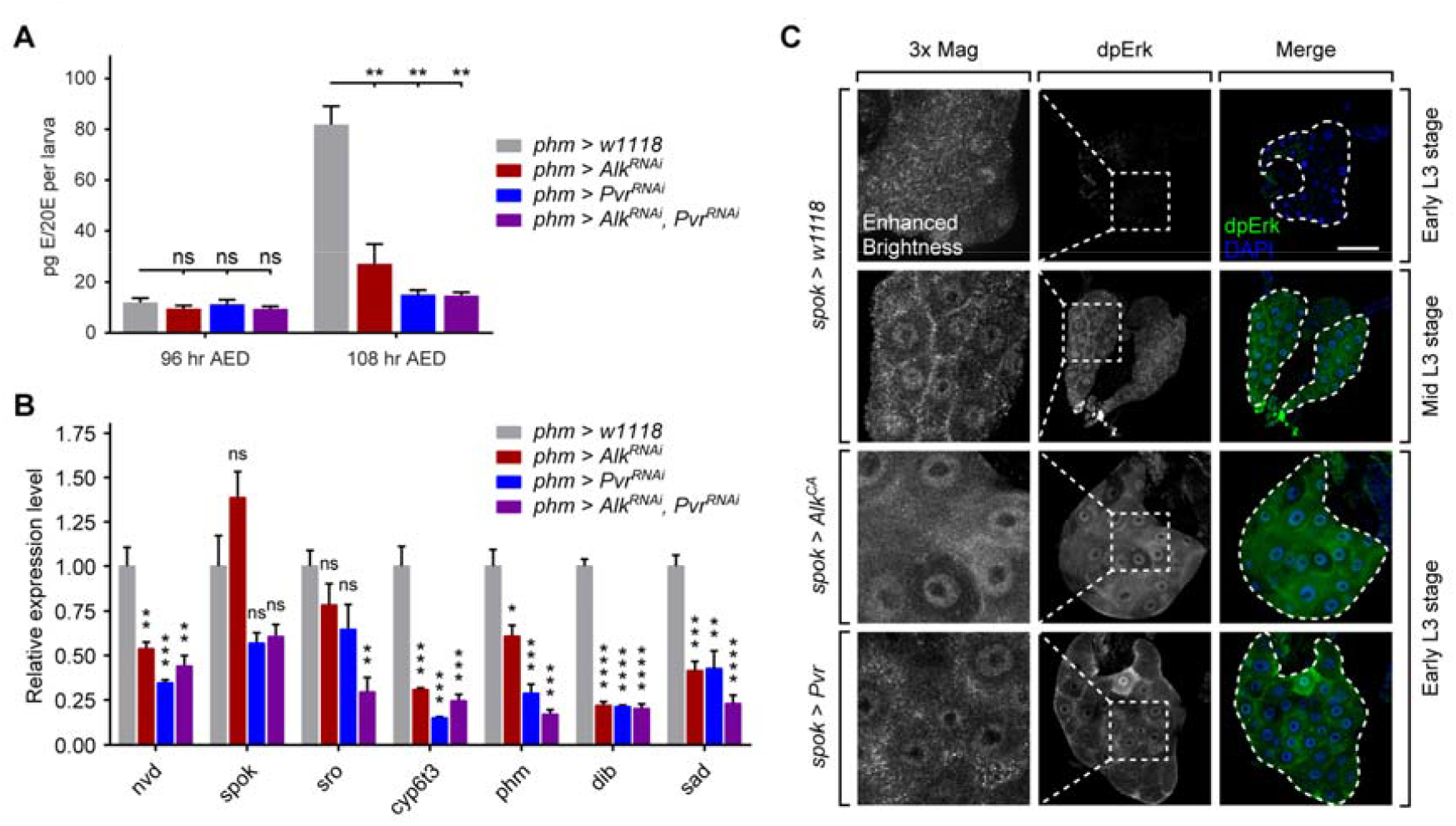
Alk and Pvr facilitate ecdysone synthesis and Halloween gene expression by activating Ras/Erk pathway. (A) Quantification of ecdysone/20-hydroxyecdysone titers in *phm*>*w1118, phm*>*Alk*^*RNAi*^, *phm*>*Pvr*^*RNAi*^ and *phm*>*Alk*^*RNAi*^, *Pvr*^*RNAi*^ larvae at indicated timing stages. (B) qRT-PCR measurements of Halloween gene expressions in wandering *phm*>*w1118* and age matched *phm*>*Alk*^*RNAi*^, *phm*>*Pvr*^*RNAi*^ and *phm*>*Alk*^*RNAi*^, *Pvr*^*RNAi*^ larvae. (A and B) Mean ± SEM; p values by unpaired t-test (n=3 in A, n=4 in B; ns, not significant, *p<0.05, **p<0.01, ***p<0.001, ****p<0.0001). (C) Immunofluorescence images of *spok*>*w1118, spok*>*Alk*^*CA*^ and *spok*>*Pvr* PGs stained with anti-phospho-Erk antibody. Dash lines outline the PG area of the ring gland in the right lane. Enlarged images of the indicated areas are shown in the left lane. To visualize the distribution of fluorescence signal in early-L3 *spok*>*w1118* larvae, the brightness is enhanced in the enlarged image. Scale bar, 50μm.

Ecdysone is synthesized from cholesterol through the action of ecdysone biosynthetic enzymes encoded by the Halloween genes (Niwa and Niwa, 2014). To determine how Alk and Pvr affects ecdysone synthesis in the PG, we assessed Halloween gene expression in *phm*>*w1118* larvae at the wandering stage and receptor suppression larvae of equivalent age. In *phm*>*Alk*^*RNAi*^, *phm*>*Pvr*^*RNAi*^ and *phm*>*Alk*^*RNAi*^, *Pvr*^*RNAi*^ larvae the expression of five out of seven Halloween genes (*nvd, cyp6t3, phm, dib, sad*) were significantly lower than *phm*>*w1118* control (Figure 2B). In addition, the expression of *sro* was suppressed in *phm*>*Alk*^*RNAi*^, *Pvr*^*RNAi*^ double suppression larvae, although in *phm*>*Alk*^*RNAi*^ or *phm*>*Pvr*^*RNAi*^ larvae no significant change was observed compared with *phm*>*w1118* (Figure 2B). These results show that Alk and Pvr signaling regulates ecdysone biosynthesis by affecting Halloween gene expression.

Previous work has established that both Alk (Englund et al., 2003; Gouzi et al., 2011; Loren et al., 2001) and Pvr (Learte et al., 2008; Sansone et al., 2015) are able to activate the Ras/Erk pathway in certain *Drosophila* embryonic and post-embryonic tissues. Thus, we tested whether the two pathways activate Ras/Erk signaling in the PG. Since other RTKs, including Torso and Egfr, also activate Ras/Erk signaling in the PG (Cruz et al., 2020; Rewitz et al., 2009), we speculated that partial suppression of Ras/Erk signaling, if it occurs, could be difficult to detect. To circumvent this possible obstacle, we asked if we could detect a change on Ras/Erk signaling in Alk and Pvr activation larvae. Unexpectedly, overexpressing Alk or Pvr using *phm-Gal4* driver caused developmental arrest at an early stage (see below), so we employed *spok-Gal4*, a weaker PG driver for receptor activation/overexpression conditions. To detect the activation level of Ras/Erk signaling, we examined PG immunofluorescence using an antibody that specifically recognizes phospho-Erk (Cruz et al., 2020; Ohhara et al., 2015). In *spok*>*w1118* larvae, the Ras/Erk signaling in the PG appears weak in the early-L3 stage and is then activated in the mid-L3 stage, as indicated by the enhanced overall immunofluorescence signal strength as well as the partial nuclear localization of the signal (Figure 2C). When constitutively activated (CA) Alk or wild type Pvr was expressed in the PG by *spok*>*Alk*^*CA*^ and *spok*>*Pvr*, respectively, Ras/Erk was strongly activated in the early-L3 stage (Figure 2C), indicating that both Alk and Pvr pathways activate Ras/Erk signaling in the PG. This result is consistent with at least partial overlap between Alk, Pvr and Torso signaling through Ras/ERK activation.

### Alk regulates autophagy in the PG by activating Pi3K/Akt pathway

In addition to Ras/Erk, Pi3K/Akt is another signaling pathway activated by RTKs (Mele and Johnson, 2019). A well-studied RTK that activates Pi3K/Akt signaling in the PG is InR, which conveys nutritional signal to the PG and promotes PG growth (Colombani et al., 2005; Mirth et al., 2005). Interestingly, one study indicates that Alk is also capable to activate Pi3K/Akt signaling and to compensate the loss of InR pathway in multiple larval tissues (Cheng et al., 2011). Therefore, we tested whether Alk and Pvr can activate Pi3K/Akt signaling in the PG. To monitor the activation of Pi3K/Akt signaling, we expressed a GFP tagged PH domain (tGPH) which binds specifically to phosphatidylinositol 3,4,5-trisphosphate (PIP3) produced by activated Pi3K. Basal level of Pi3K/Akt activation was observed in the PGs of *spok*>*w1118* larvae, indicated by the GFP signal on PG cell membrane (Figure 3A). The membrane localizes GFP signal was much stronger in the PGs expressing activated InR, consistent with the known capability of InR to activate Pi3K/Akt signaling (Weinkove and Leevers, 2000). Comparable strong membrane GFP signal was observed in *spok*>*Alk*^*CA*^ PG cells (Figure 3A), showing that Alk activation can induce Pi3K/Akt signaling in the PG. However, no such signal was identified in *spok*>*Pvr* PGs (Figure 3A). Since both InR and Alk activate Pi3K/Akt signaling, we sought to determine whether Alk can compensate the loss of InR signaling in the PG. Overexpressing either activated InR or Alk caused precocious pupariation (Figure 3B), suggesting similar activities of the two receptors in the PG. Suppressing InR activity by *phm*>*InR*^*DN*^ delayed the timing of pupariation, which is effectively rescued by activated Alk (Figure 3B). These results suggest that Alk activates Pi3K/Akt signaling and perhaps functions to supplement the InR pathway in the PG.

**Figure 3.**
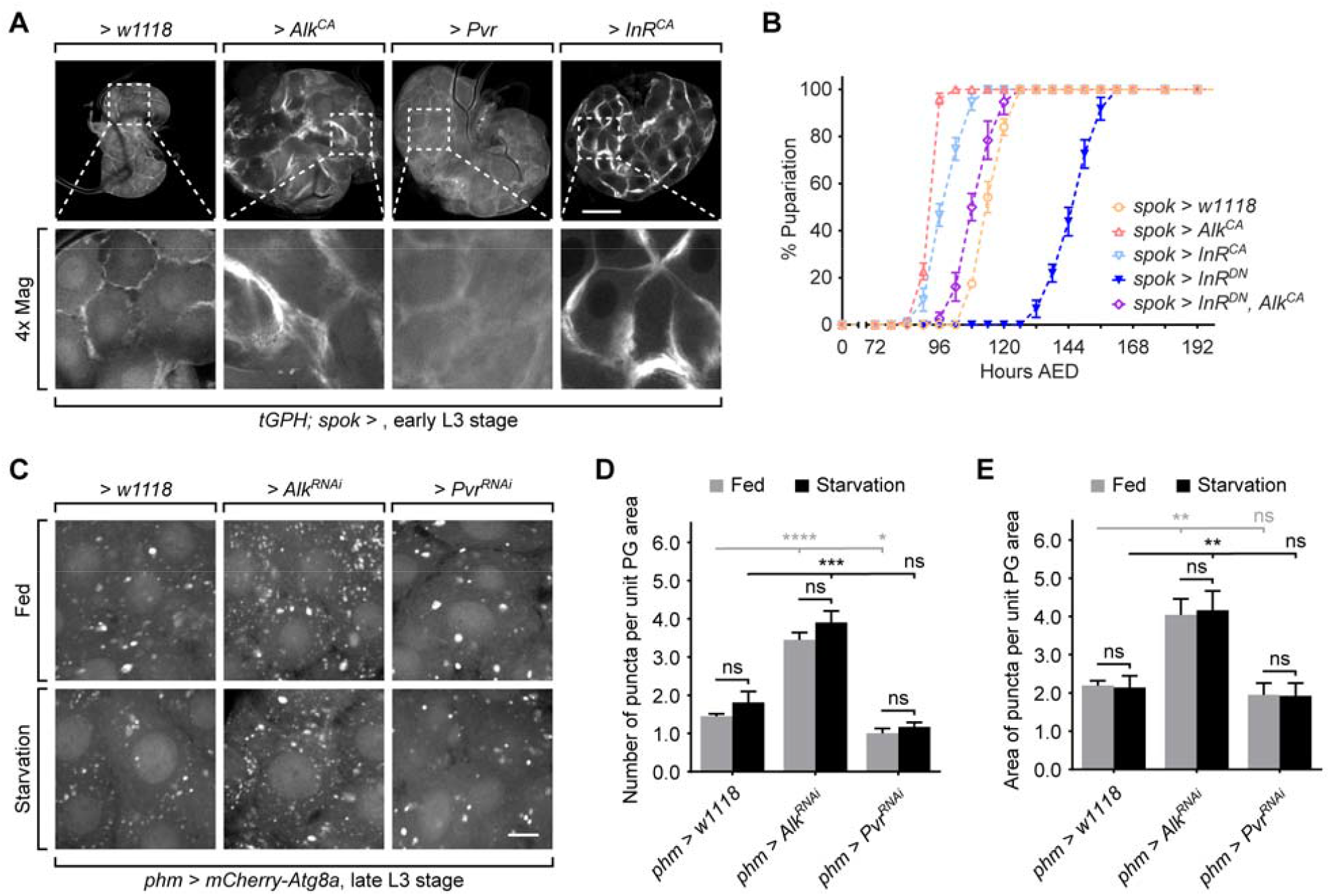
Alk regulates autophagy in the PG by activating Pi3K/Akt pathway. (A) Images of *spok*>*w1118, spok*>*Alk*^*CA*^, *spok*>*Pvr* and *spok*>*InR*^*CA*^ PGs expressing *tGPH*. Enlarged images of indicated areas are also shown. Scale bar, 50μm. (B) Pupariation timing curves of *spok*>*w1118, spok*>*Alk*^*CA*^, *spok*>*InR*^*CA*^, *spok*>*InR*^*DN*^ and *spok*>*InR*^*DN*^, *Alk*^*CA*^ larvae. (C) Images of *phm*>*w1118, phm*>*Alk*^*RNAi*^ and *phm*>*Pvr*^*RNAi*^ PGs expressing mCherry-Atg8a. Animals were starved at late-L3 stage for 4 hours to induce autophagy. Scale bar, 10μm. (D and E) Quantification of the number (D) and the total area (E) of Atg8a-positive puncta per unit PG cell area. Mean ± SEM; p values by unpaired t-test (n=5-7; ns, not significant, *p<0.05, **p<0.01, ***p<0.001, ****p<0.0001).

Autophagy is a process modulated by Pi3K/Akt signaling that has been reported to regulate ecdysone biosynthesis by altering cholesterol metabolism in the PG (Pan et al., 2019; Texada et al., 2019). Thus we tested whether Alk suppression affects autophagy induction in the PG. Previously, we have shown that autophagy is strongly induced by starvation in the early-, but not the late-L3 stage (Pan et al., 2019). Since Alk is highly expressed in the late-L3 stage (Figure 1B), we hypothesized that Alk signaling may be responsible for suppressing of autophagy induction during late stage development. To test this possibility, we analyized autophagy induction in the PG using *phm*>*mCherry-Atg8a* in fed and starved late-L3 larvae. By measuring both the number and the area of Atg8a-positive puncta, we found that autophagy is significantly induced in *phm*>*Alk*^*RNAi*^ larvae in both fed and starvation condition (Figure 3C-E). In contrast, knocking down Pvr in the PG did not affect autophagy induction (Figure 3C-E), in agreement with the finding that Pvr does not induce Pi3K/Akt signaling. Taken together, we conclude that Alk, but not Pvr, suppresses the inducibility of PG autophagy induction in late stage larvae.

### Activation of Alk and Pvr pathway affects developmental timing in a dose dependent manner

Since suppression of Alk and Pvr delays the timing of pupariation, we tested whether activation of the receptors accelerates developmental timing. Alk and Pvr activation by *spok*>*Alk*^*CA*^ and *spok*>*Pvr* resulted in earlier pupariation and formation of smaller pupae (Figure 4A-B), which is consistent with our hypothesis that they contribute to the developmental timing signal. Upon activation of Alk or Pvr in *ptth* mutants, the developmental delay and larger pupal size caused by loss of *ptth* was reversed by Alk and Pvr activation (Figure 4C-D), showing that activation of Alk and Pvr can compensate for loss of PTTH/Torso signaling.

**Figure 4.**
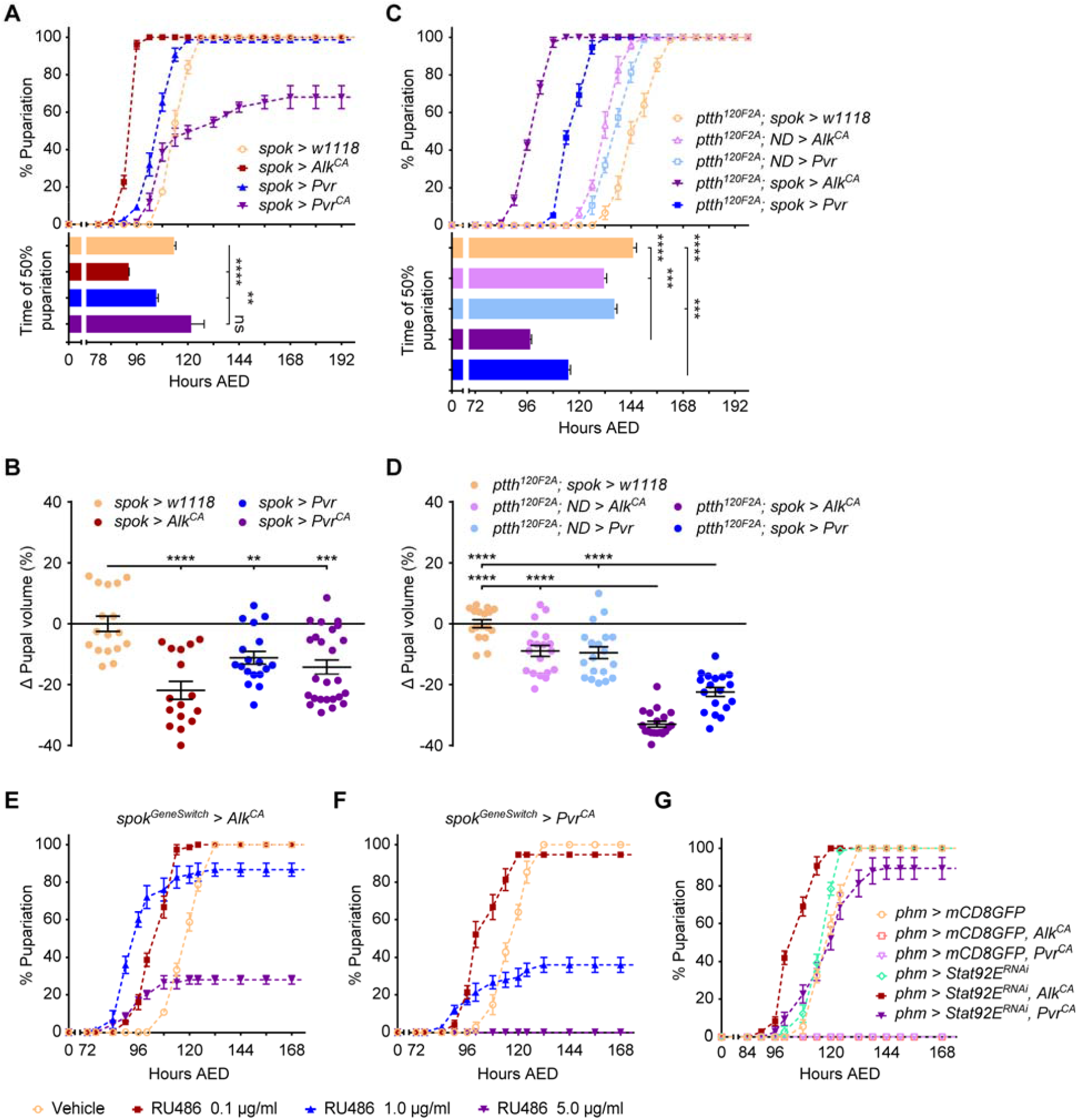
Activation of Alk and Pvr pathway affects developmental timing in a dose dependent manner. (A) Pupariation timing curves and the time of 50% pupariation of *spok*>*w1118, spok*>*Alk*^*CA*^, *spok*>*Pvr* and *spok*>*Pvr*^*CA*^ larvae. (B) Relevant pupal volume changes in animals tested in (A). (C) Pupariation timing curves and the time of 50% pupariation of *spok*>*w1118, spok*>*Alk*^*CA*^ and *spok*>*Pvr* larvae with *ptth*^*120F2A*^ mutant background. ND, no driver. (D) Relevant pupal volume changes in animals tested in (C). (E and F) Pupariation timing curves of *spok*^*GeneSwitch*^>*Alk*^*CA*^ (E) and *spok*^*GeneSwitch*^>*Pvr*^*CA*^ (F) larvae fed with indicated concentrations of RU486. (G) Pupariation timing curves of *phm*>*w1118, phm*>*Alk*^*CA*^ and *phm*>*Pvr*^*CA*^ larvae with/without knockdown of Stat92E. To balance the driver strength of phm-Gal4, *UAS-mCD8GFP* was introduced to the groups without *UAS-Stat92E*^*RNAi*^. (A-D) Mean ± SEM; p values by unpaired t-test (n=3 in A and C, n=16-25 in B and D; ns, not significant, **p<0.01, ***p<0.001, ****p<0.0001).

Curiously, when Pvr signaling is further increased through expression of constitutively activated Pvr (*spok*>*Pvr*^*CA*^) many larvae failed to pupariate and the rest pupariated no early than the *spok*>*w1118* controls (Figure 4A). Furthermore, as mentioned above, overexpressing Alk and Pvr using the strong *phm-Gal4* PG driver results in developmental arrest before larvae reach L3 stage. Based on these observations using different Alk/Pvr activation models, we hypothesized that the effect of Alk and Pvr activation on developmental timing is “dose dependent”. That is, weak/moderate activation of Alk and Pvr causes precocious pupariation, but high-level activation leads to detrimental effects on development. To verify this “dose dependence” hypothesis, we employed a *spok*^*Geneswitch*^*-Gal4* whose Gal4 driver strength is determined by the concentration of RU486 administration (Roman et al., 2001). In *spok*^*GeneSwitch*^>*Alk*^*CA*^ larvae, a low dose of RU486 feeding (0.1 μg/ml) caused earlier pupariation, while a high dose (5.0 μg/ml) led to a high rate of developmental arrest in L3 stage (Figure 4E). A mid-level dose (1.0 μg/ml) caused a mixed phenotype of precocious pupariation and developmental arrest (Figure 4E), confirming the bi-phased, “dose dependent” effects of Alk activation of developmental timing. Similar results were observed in *spok*^*GeneSwitch*^>*Pvr*^*CA*^ larvae, the only difference being that the medium dose RU486 caused a higher rate of developmental arrest (Figure 4F). These data demonstrate that moderate, but not high-level activation of Alk and Pvr accelerates the timing of pupariation.

To explore the mechanism underlying the detrimental effect caused by receptor overactivation, we initially examined PG morphology in the receptor activation larvae. PG tissue overgrowth was found in all *spok*>*Alk*^*CA*^, *spok*>*Pvr* and *spok*>*Pvr*^*CA*^ larvae (Figure S2A). However, only the *spok*>*Pvr* PG exhibited uniform cell and nucleus sizes, which is also observed in *spok*>*InR*^*CA*^ PGs (Figure S2A). In both *spok*>*Alk*^*CA*^ and *spok*>*Pvr*^*CA*^ PGs, cells exhibited intensive heterogeneity and loss of normal tissue organization (Figure S2A), reminiscent of the atypical morphology of cancerous tissues. Based on these observations, we speculate that the atypical growth of PG is likely even worse in *phm-Gal4* driven receptor overactivation animals and it results in developmental arrest as a result of PG cell malfunction or even death.

Previous studies have shown that simultaneous activation of Ras/Erk and Jak/Stat signaling induces cancerous-like growth in *Drosophila* larval tissues (Wu et al., 2010). Gain-of-function alleles of Torso has also been found to induce activation of Jak/Stat pathway during embryonic development (Li et al., 2002). Inspired by these observations, we tested whether Jak/Stat signaling is activated by either Alk or Pvr. Using 10xStat92E-GFP, a reporter of Jak/Stat signaling, we observed remarkably strong GFP signal in *spok*>*Alk*^*CA*^, *spok*>*Pvr* and *spok*>*Pvr*^*CA*^ PGs (Figure S2B), clearly showing that both Alk and Pvr activation can induce Jak/Stat signaling in the PG. Interestingly, *spok*>*Torso* did not induce strong Jak/Stat activation in the PG, despite the ability of activated alleles to do so in some embryonic tissues (Figure S2B) (Li et al., 2002), perhaps again indicating that dose/strength is an important factor to consider when considering which downstream pathways can be activated by these different RTKs

We next investigated whether Jak/Stat signaling mediated the developmental defects caused by Alk and Pvr overactivation. We used *phm-Gal4* to induce overactivation of the receptors and suppressed the Jak/Stat pathway by *UAS-Stat92E*^*RNAi*^. Since *UAS-Stat92E*^*RNAi*^ could weaken the driver strength of *phm-Gal4*, we introduced *UAS-mCD8GFP* in the control groups without *UAS-Stat92E*^*RNAi*^ to equalize the UAS transgene copy number. Both *phm*>*mCD8GFP, Alk*^*CA*^ and *phm*>*mCD8GFP, Pvr*^*CA*^ larvae arrested at various larval stages before pupariation (Figure 4G). Knockdown of *Stat92E* did not significantly affect developmental timing by itself. However, the developmental arrest caused by *phm*>*mCD8GFP, Alk*^*CA*^ and *phm*>*mCD8GFP, Pvr*^*CA*^ were effectively rescued in *phm*>*Stat92E*^*RNAi*^, *Alk*^*CA*^ and *phm*>*Stat92E*^*RNAi*^, *Pvr*^*CA*^ animals, respectively (Figure 4G). These results demonstrate that Jak/Stat signaling induced by Alk and Pvr overactivation mediates the developmental defects in Alk and Pvr overactivation animals. Since Jak/Stat is very weakly induced in *phm*>*w1118* control animals (Figure S2B), we conclude that Alk and Pvr do not strongly activate Jak/Stat signaling in wild type animals but may do so under certain developmental or environmental conditions.

### Ligands that activate Alk and Pvr derive from both PGNs and PG

After confirming the effect of Alk and Pvr receptors on developmental timing and body size control, we sought to determine the source of their ligands that activate the receptors in the PG. Based on our previous observations that ablation of PGNs produces a stronger phenotype than loss of *ptth* (Shimell et al., 2018), we speculate that some proportion the ligands may derive from the PGNs. However, autocrinal regulation pathways have also been discovered in the PG (Cruz et al., 2020; Ohhara et al., 2015), indicating that the ligands may also be produced by the PG itself. Therefore, we tested for ligand expression in both PGs and PGNs.

Jelly belly (Jeb) has been identified as the only known ligand for Alk (Englund et al., 2003). To examine the expression pattern of Jeb, we took advantage of the Minos Mediated Integration Cassette (MiMIC) insertion fly line (*Jeb*^*MI03124*^) (Nagarkar-Jaiswal et al., 2015) and converted it into a Gal4 expression line (*Jeb*^*T2A*^*-Gal4*) using Recombinase-Mediated Cassette Exchange (RMCE) strategy (Diao et al., 2015). In the *Jeb*^*T2A*^>*EGFP* larvae, we observed Jeb expression in a number of cells in larval brain lobes, while no obvious expression was detected in the PG (Figure S3A). By immunostaining using anti-PTTH antibody, we clearly found overlap between the EGFP and the PTTH signals (Figure 5A), showing that Jeb is expressed in the PTTH producing neurons.

**Figure 5.**
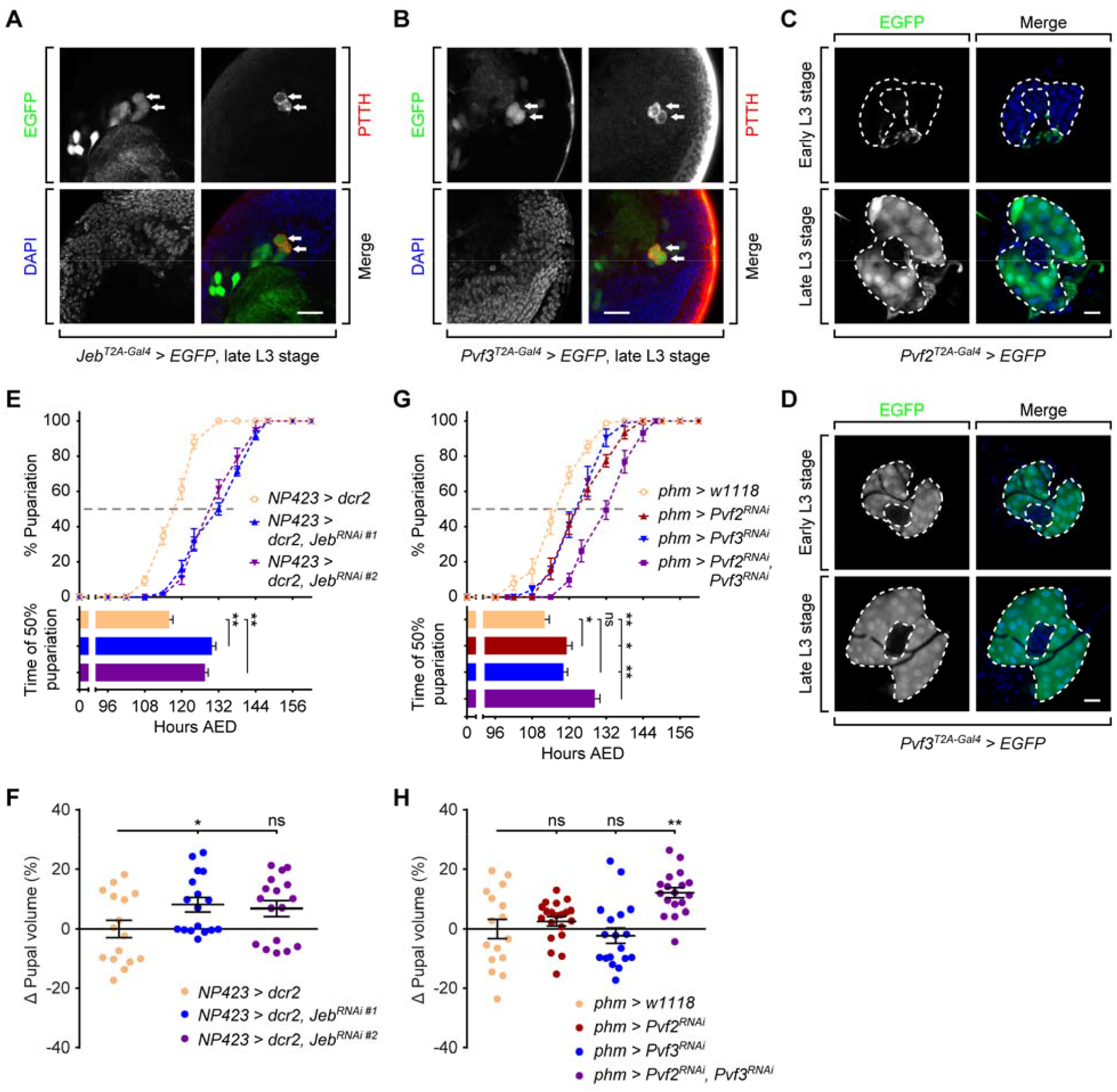
Ligands that activate Alk and Pvr derive from both PGNs and PG. (A and B) Immunofluorescence images of *Jeb*^*T2A-Gal4*^>*EGFP* (A) and *Pvf3*^*T2A-Gal4*^>*EGFP* (B) larval brains stained with anti-PTTH antibody. Arrows indicate the co-localization between the EGFP and PTTH immunofluorescence signals. Scale bar, 20μm. (C and D) Images of *Pvf2*^*T2A-Gal4*^>*EGFP* (A) and *Pvf3*^*T2A-Gal4*^>*EGFP* (B) PGs which has expression of EGFP in Pvf2- and Pvf3-expressing cells. Dash line marks the outline of PG area in the ring gland. Scale bar, 50μm. (E) Pupariation timing curves and the time of 50% pupariation of *NP423*>*w1118* and two groups of *NP423*>*Jeb*^*RNAi*^ larvae. (F) Relevant pupal volume changes in animals tested in (E). (G) Pupariation timing curves and the time of 50% pupariation of *phm*>*w1118, phm*>*Pvf2*^*RNAi*^, *phm*>*Pvf3*^*RNAi*^ and *phm*>*Pvf2*^*RNAi*^, *Pvf3*^*RNAi*^ larvae. (H) Relevant pupal volume changes in animals tested in (G). (E-H) Mean ± SEM; p values by unpaired t-test (n=3 in E and G, n=16-20 in F and H; ns, not significant, *p<0.05, **p<0.01).

Unlike Alk, Pvr has three known ligands - Pvf1, Pvf2 and Pvf3. To determine which of the Pvf ligands activate Pvr in the PG, we first tested the viability of null mutants of the three genes. The *Pvf2*^*T2A*^*-Gal4* and *Pvf3*^*T2A*^*-Gal4* larvae, in which the endogenous *Pvf* gene expression is abrogated by the T2A cassette insertion (Diao et al., 2015), did not survive into L3 stage. However, a well characterized null mutant *Pvf1*^*EP1624*^ (Duchek et al., 2001) pupariated without significant delay (Figure S3B). Therefore, we propose that Pvf2 and Pvf3 may be the ligands that interact with Pvr in the PG. Using the T2A-Gal4 lines, we observed Pvf3 expression in the PGNs (Figure 5B), while both Pvf2 and Pvf3 are expressed in the PG (Figure 5C and D). Intriguingly, the expression of Pvf2 and Pvf3 in the PG exhibited different temporal patterns. Pvf2 expression is limited in early-L3 stage but surges in late-L3 stage (Figure 5C), while Pvf3 expression is kept at high level throughout the L3 stage (Figure 5D).

We next tested whether these ligands are required for the effects of Alk and Pvr signaling on pupariation timing control. Using the *NP423-Gal4* driver, we first knocked down Jeb or Pvf3 in the PGNs (Yamanaka et al., 2013b). Depletion of Jeb in the PGNs using two RNAi constructs caused delayed pupariation and enlarged pupal size (Figure 5E and F), showing that the activation of Alk in the PG is, at least partially, mediated from the PGN-derived Jeb signal. However, knockdown of Pvf3 in the PGNs did not significantly affect timing of pupariation (Figure S3C and D), indicating that the Pvr signaling does not fully rely on the PGN-derived Pvf3. Next we suppressed the expression of Pvf2 and Pvf3 in the PG using multiple RNAi constructs. Neither Pvf2 nor Pvf3 knockdown caused a significant delay in developmental timing, except for one Pvf2 RNAi (*phm*>*Pvf2*^*RNAi #3*^) which resulted in a minor timing delay (Figure S3E-H). However, when both ligands were simultaneously knocked down in the PG using *phm*>*Pvf2*^*RNAi*^, *Pvf3*^*RNAi*^, we observed significant delay of timing and enlarged pupae compared with *phm*>*w1118* control (Figure 5G and H). This result shows that both Pvf2 and Pvf3 likely activate Pvr following an autocrine pathway, although some contribution of PGN-derived Pvf3 cannot be ruled out.

Lastly, we examined whether overexpression of the ligands could phenocopy the receptor activation animals. Neither Jeb and Pvf3 overexpression in the PGNs nor Pvf2 and Pvf3 overexpression in the PG induced a significant change on the timing of pupariation (Figure S3I-L), showing that the activity of Alk and Pvr pathways in the PG is not solely controlled by expression of the ligands.

## Discussion

### Multiple RTK signals coordinate in the PG to regulate developmental timing

In previous studies, three RTKs, Torso (Rewitz et al., 2009), InR (Colombani et al., 2005; Mirth et al., 2005) and Egfr (Cruz et al., 2020), have been proved crucial in the PG for the control of pupariation and body size. In this work, we identified two additional RTKs, Alk and Pvr, that are also required for proper timing and body size control. Suppression of either Alk or Pvr compromises E synthesis in the PG (Figure 2A), delays pupariation (Figure 1A) and increases pupal size (Figure 1C), while moderate activation of Alk or Pvr accelerates development (Figure 4A). The biological functions of Alk/Pvr in the neuroendocrine pathway are similar to those of the other RTKs (Colombani et al., 2005; Cruz et al., 2020; Mirth et al., 2005; Rewitz et al., 2009), indicating likely signal coordination among the receptors. Downstream of the receptors, Torso (Rewitz et al., 2009), Egfr (Cruz et al., 2020), Alk and Pvr (Figure 2C) Ras/Erk signaling is activated, while InR (Mirth et al., 2005) and Alk (Figure 3A) stimulate the Pi3K/Akt pathway. Consistent with signal pathway convergence, suppression of Alk and Pvr simultaneously or suppression of Alk/Pvr in *ptth* mutants exhibit prolonged delay of developmental timing and larger pupal size (Figure 1A and C-E). In addition, activation of Alk/Pvr rescues the developmental defects in *ptth* mutants (Figure 4C and D), while activated Alk antagonizes the suppression of InR (Figure 3B). In total, both the downstream signaling pathway convergence and the additive effects of receptor activation/suppression support the coordination of signaling among these RTKs. Cellular level coordination of receptor-mediated signals is very common during development. The PG is a good example of this coordination, as it integrates a large variety of signals, such as PTTH (Shimell et al., 2018), Hedgehog (Palm et al., 2013; Rodenfels et al., 2014), Activin (Gibbens et al., 2011), Bone Morphogenetic Protein (BMP) (Setiawan et al., 2018), etc., and interprets them to produce a precisely controlled amount of hormone. At least four RTKs (Torso, Egfr, Alk and Pvr) are expressed in the PG, all of which activate the Ras/Erk pathway (Cruz et al., 2020; Rewitz et al., 2009) (Figure 1B and 2C). Although PTTH/Torso has been considered the key tropic signal for PG function, actually all three of the other RTKs can take the place of Torso to maintain some level of PG ecdysone production (Cruz et al., 2020) (Figure 1D and E). Loss of either Torso, Alk or Pvr signal causes developmental delay but does not block pupariation (Figure 1A and D). Even considering that loss of Egfr in the PG causes arrest at the L3 stage (Cruz et al., 2020), Egfr is still dispensable during the first two molts which also require production of E pulses by the PG. These observations lead to an open question: why does the PG utilize multiple signals that appear to function redundantly?

An obvious possibility is that multiple timing signals provide both robustness and flexibility in response to variable developmental conditions. For example, given a choice of diets *Drosophila* larva chose one that maximizes developmental speed over other life-history traits (Rodrigues et al., 2015). This is not surprising given the ephemeral nature of rotting fruit, a primary food source for *Drosophila*. Thus, multiple signals may enable larva to maximize developmental speed. Another possibility is that the different signals contribute to different temporal aspects of the developmental profile. For example, perhaps none of the receptors alone can achieve a strong enough Ras/Erk activation in late stage larva that meets the demand for large rise in E production that triggers wandering and initiation of pupation. Interestingly, the expression of Egfr (Cruz et al., 2020), Alk and Pvr (Figure 1B) all increase remarkably during the late L3 stage when both Halloween gene expression and E synthesis ramps up, suggesting that the three receptors may function as supplements to Torso in order to achieve robust Ras/Erk activation and stimulation of ecdysone production.

Yet another possibility is that in addition to Ras/Erk signaling, each receptor may induce other downstream pathway(s). For instance, we have previously reported that regulated autophagy induction in the PG is a key mechanism that prevents precocious non-productive pupation by limiting E availability if larva have not achieved CW (Pan et al., 2019). In that report, we also demonstrated that after CW autophagy inducibility is greatly repressed. This makes sense from a developmental perspective because if food becomes limiting after CW is achieved, it would seem disadvantageous to slow development down by limiting E production. Therefore a mechanism to shut down autophagy inducibility after attainment of CW would appear to be beneficial and as reported here we find that Alk activation is, in part, responsible for shut down of autophagy activation in the PG after the CW nutrient checkpoint has been surpassed (Figure 3C-E). Lastly, since Drosophila is a highly derived insect some of these signals may represent more basal or ancient mechanisms for triggering E pulses. For example, PTTH and/or Torso are missing in several insect species including, cockroaches, honeybees and some parasitic wasps (Skelly et al. 2018) yet these species are able to undergo metamorphosis and may employ some combination of these other signals as timing cues.

### Activation of Alk/Pvr pathways results in dose dependent effects on development via Jak/Stat signaling

Our manipulations of Alk and Pvr but not Torso signaling in the PG lead to the discovery that Jak/Stat activation can also affect developmental timing (Figure S2B). A distinct feature of Alk and Pvr is that they can exert opposite effects on development likely depending on the activation strength. Weak activation of Alk or Pvr in the PG facilitates pupariation, while strong activation results in arrest of development at various larval stages (Figure 4E and F) due to Jak/Stat activation. Using weak *spok-Gal4* driver leads to overgrowth of the PG and to atypical morphology (Figure S2A). Tissue overgrowth is commonly observed when either Pi3K/Akt or Ras/Erk is hyperactivated in the PG, however, neither pathway induces atypical morphological change in the overgrown PGs or developmental arrest (Caldwell et al., 2005; Mirth et al., 2005) as we observe when Alk or Pvr are hyperactivated especially with the strong *phm-Gal4* driver. Since suppression of Jak/Stat rescues the developmental arrest caused by *phm-Gal4* driven Alk/Pvr hyperactivation (Figure 4G), it appears that Jak/Stat signaling is the key factor that mediates the side effect of Alk/Pvr activation on PG morphology and developmental timing. At lower levels of activation as found in the *spok*>*Alk*^*CA*^ and *spok*>*Pvr*^*CA*^ many larvae still manage to pupariate (Figure 4A), suggesting that larvae can tolerate a certain level of ectopic Jak/Stat signaling caused by Alk/Pvr activation. What goes wrong at high level activation of Jak/Stat is still not clear nor do we know what the endogenous late Jak/Stat signal contributes in terms of PG function since knockdown with available reagents did not produce a significant delay phenotype (Figure 4G). In *Drosophila*, canonical Jak/Stat signaling pathway is commonly induced by a group of cytokines including Unpaired 1-3 (Upd1-3) via their cognate receptor Domeless (Dome) (Trivedi and Starz-Gaiano, 2018). However, it has also been reported that Torso and Pvr are capable of inducing Jak/Stat activation in some circumstances (Li et al., 2002; Mondal et al., 2011). Although we did not observe induction of Jak/Stat signal by overexpressing wildtype Torso in the PG (Figure S2B), this might be due to a weaker activation using wildtype Torso overexpression versus gain-of-function *tor*^*Y9*^ and *tor*^*RL3*^ mutants as used in the previous study (Li et al., 2002). Since we observe Dome expression and endogenous activation of the 10xStat92E-GFP reporter in late-L3 PGs (Figure S2C), we assume it is likely to play some role at this stage. Whether the Jak/Stat activation is through Alk/Pvr or via reception of canonical Upd/Domeless signals is not clear. It is interesting to note that Upd2 is secreted from the fat body into hemolymph (Rajan and Perrimon, 2012) and therefore may provide a nutrient storage signal to the PG that could be important regulator of developmental timing perhaps under certain types of non-standard lab growth conditions, while Upd3 is produced by hemocytes and affects multiple tissue and could thereby provide an immune response signal that could also modulate developmental timing (Shin, et al. 2020).

### Ligands activate Alk/Pvr through both neuronal and autocrine pathways

Since its discovery, PTTH has been recognized as the most important prothoracicotropic neuropeptide that triggers metamorphosis in holometabolous insects (McBrayer et al., 2007; Shimell et al., 2018; Smith and Rybczynski, 2012). In some species, such as *Bombyx mori*, additional prothoracicotropic neuropeptides such as Orcokinin (Yamanaka et al., 2011) and FXPRL-amide peptides (Watanabe et al., 2007) have been discovered, however, PTTH and insulin like peptides (Ilps) are the only known brain-derived PG tropic hormones in *Drosophila*. Nevertheless, analysis of the Drosophila PTTH null mutant phenotype verses PGN ablation and PGN electrical manipulation provided evidence that there are other tropic signals derived from the Drosophila PGNs (McBrayer et al., 2007; Shimell et al., 2018). Our observations described here demonstrate that the Alk ligand Jeb and the Pvr ligand Pvf3 are produced in the PGNs (Figure 5A and B). Knockdown of Jeb in the PGNs causes delay of pupariation and increased pupal size (Figure 5E and F), phenocopying the *phm*>*Alk*^*RNAi*^ animals (Figure 1A and C) and showing that the PGNs are the major source of Jeb that functions in the PG. Depletion of Pvf3 in the PGNs does not significantly affect developmental timing (Figure S3C and D), which is not a surprise since we found that Pvf2 and Pvf3 are also produced in the PG itself (Figure 5C and D). Overexpressing neither Jeb nor Pvf3 in the PGNs was found to influence timing (Figure S3I and J), indicating that the neural activity of PGNs and/or the temporal regulation of Alk/Pvr expression plays the dominant role in the regulation of signaling by these factors.

Besides the well-established role of the PGNs in regulating developmental timing and body size, several recent studies also indicate that autocrine signaling within the PG itself provide important developmental regulatory cues. This was first documented for biogenic amine signaling (Ohhara et al., 2015) but more recently extended to include the RTK Egfr and its ligands Vein and Spitz (Cruz et al., 2020). Interestingly, the expression of Vein and Spitz in the PG increase in mid to late L3 and may not contribute to CW determination, but instead respond to it to form part of a E feedforward circuit that helps ramp up hormone production during late L3 in anticipation of the large pulse that drives pupation (Cruz et al., 2020; Moeller et al., 2013). Similarly, since we observe expression of both Pvf2 and Pvf3 in the late L3 PG (Figure 5C and D) and since knockdown of Pvf2 and Pvf3 simultaneously in the PG causes delay of pupariation and larger pupal size (Figure 5G and H), these ligands together with their receptor Pvr also appear to form a autocrine signaling pathway. We and others have also observed expression of Pvf2/3 in other tissues/cell types such as fat body, salivary gland (data not shown) and hemocytes (Parsons and Foley, 2013). Whether these sources also provide some input to the PG is not clear. We also found that overexpression of neither Pvf2 nor Pvf3 caused accelerated development (Figure S3K and L), which is in stark contrast to the case of Egfr signaling in which overexpression of Vein or Spitz advances pupariation significantly (Cruz et al., 2020). This finding indicates that the activity of Pvr signaling may depend on the expression of Pvr receptor and/or the release of ligands, rather than ligand expression. Endogenous Pvf2 expression is limited to the late-L3 stage, yet Pvf3 is constitutively expressed in the L3 stage (Figure 5C and D). The biological significance of the differentially regulated Pvf ligand expression is still an open question. It is noteworthy that there are three *Pvr* isoforms produced by alternative splicing among the exons coding the ligand binding domain (Cho et al., 2002; Hoch and Soriano, 2003). Thus, reception of different Pvf ligand signals could very much depend on the levels and timing of receptor isoform expression in the PG. Lastly, we note that neither Alk nor Pvr accumulate to substantial levels on the PG membrane until after CW (Figure 1B). Thus, like Egfr signaling, their primary functions likely control post-CW events. What regulates the post-CW membrane localization of these receptors is not yet clear, but it is interesting to speculate that the process might be one of the first downstream responses to surpassing the CW checkpoint that prepares the PG gland for a major acceleration in hormone production.

## Acknowledgements

We thank Dr. Ruth Palmer, Dr. Edan Foley, Dr. Norbert Perrimon and Dr. Ben-Zion Shilo for fly line and reagent sharing. We also thank the Vienna Drosophila RNAi Center, the Bloomington Drosophila Stock Center and the National Institute of Genetics in Japan for fly stocks. This work was supported by grant 1R35GM118029 from NIGMS to M.B.O.

## Author contributions

Conceptualization, M.B.O. and X.P.; Methodology, M.B.O. and X.P.; Investigation, X.P.; Writing - Original Draft, X.P.; Writing - Review & Editing, M.B.O. and X.P.; Funding Acquisition, M.B.O.; Supervision, M.B.O.

## Materials and Methods

### Contact for Reagent and Resource Sharing

Further information and requests for resources and reagents should be directed to and will be fulfilled by the Lead Contact, Michael B. O’Connor.

## Experimental Model and Subject Details

### Flies

Unless noted, all flies were reared on standard agar-cornmeal food supplemented with yeast at 25□°C. Flies were cultured in 12:12 light-dark cycles, however, all experiments were carried out under constant light to avoid the potential impact of circadian cycle on developmental timing. *Phm-Gal4* (Ono et al., 2006) and *spok-Gal4* (Moeller et al., 2017) was used to drive gene expression specifically in PG cells. *NP423-Gal4* (Yamanaka et al., 2013b) was used to drive gene expression in the PGNs. *Dome-Gal4* (Ghiglione et al., 2002) (gift from Dr. Norbert Perrimon) was used to examine the expression of *Domeless* in the PG. *Spok-GeneSwitch-Gal4* (Zeng et al., 2020) was used to drive temporally specific gene expression under control of RU486 administration. A collection of RNAi strains from Transgenic RNAi Project (TRiP) (Ni et al., 2011) were obtained from Bloomington Stock Center (BDSC) and used to carry out the targeted screen of RTKs: *UAS-Alk-RNAi* (JF02668), *UAS-btl-RNAi* (HMS02038), *UAS-Cad96Ca-RNAi* (HMC04150), *UAS-CG10702-RNAi* (HMS02499), *UAS-Ddr-RNAi* (HMC04190), *UAS-dnt-RNAi* (HMC06353), *UAS-drl-RNAi* (HMS01918), *UAS-Drl-2-RNAi* (HMC04172), *UAS-Egfr-RNAi* (JF01368), *UAS-Eph-RNAi* (HMS01986), *UAS-htl-RNAi* (HMS01437), *UAS-InR-RNAi* (HMS03166), *UAS-Nrk-RNAi* (HMC03875), *UAS-otk-RNAi* (HMC04139), *UAS-Pvr-RNAi* (HMS01662), *UAS-Ret-RNAi* (HMC04143), *UAS-Ror-RNAi* (HMC05341), *UAS-sev-RNAi* (HMC04136), *UAS-Tie-RNAi* (HMJ21428), *UAS-tor-RNAi* (HMS00021). Additional RNAi lines were used for gene knockdown, including *UAS-Stat92E-RNAi* (HMS00035), *UAS-Jeb-RNAi* (HMC04318), *UAS-Pvf2-RNAi* (HMJ23540), *UAS-Pvf3-RNAi* (HMS01876) from TRiP, *UAS-Alk-RNAi* (v107083), *UAS-Pvr-RNAi* (v43459), *UAS-Pvr-RNAi* (v43461), *UAS-Pvr-RNAi* (v105353), *UAS-Jeb-RNAi* (v103047), *UAS-Pvf2-RNAi* (v7629), *UAS-Pvf3-RNAi* (v37933) from Vienna Drosophila Resource Center (VDRC) and *UAS-Pvf2-RNAi* (13780R-2), *UAS-Pvf3-RNAi* (13781R-1) from National Institute of Genetics (NIG), Japan. *UAS-Alk*^*CA*^ (Zettervall et al., 2004), *UAS-Alk*^*DN*^ (Bazigou et al., 2007) and *UAS-Jeb* (Varshney and Palmer, 2006) lines (gifts from Dr. Ruth Palmer) were used to manipulate Alk signaling. *UAS-Pvr* (BDSC #58998), *UAS-Pvr*^*CA*^ (BDSC #58428), *UAS-Pvr*^*DN*^ (BDSC #58431), *UAS-Pvf2* and *UAS-Pvf3* lines (gifts from Dr. Edan Foley) were used to manipulate Pvr signaling. UAS-Torso and *UAS-InR*^*CA*^ (BDSC #8440) lines were used to manipulate Torso and InR signaling, respectively. *tGPH* (BDSC #8163) and *10xStat92E-GFP* (BDSC #26197) lines were used to monitor activation of Pi3K/Akt and Jak/Stat pathway, respectively. *Jeb*^*T2A*^*-Gal4, Pvf2*^*T2A*^*-Gal4* and *Pvf3*^*T2A*^*-Gal4* lines were generated from *Jeb*^*Mi03124*^ (BDSC #36200), *Pvf2*^*MI00770*^ (BDSC #32696) and *Pvf3*^*MI04168*^ (BDSC #37270), respectively, following recombinase-mediated cassette exchange strategy (Diao et al., 2015) and were used to examine the expression pattern of the corresponding genes. *Ptth*^*120F2A*^ (Shimell et al., 2018)and *Pvf1*^*EP1624*^ (BDSC #11450) null mutant lines were also used in the study.

## Method details

### Developmental timing measurement

Before egg collection, flies were transferred to constant light environment for at least 2 days and all subsequent treatments were carried out under constant light. Eggs were collected on apple juice plates with yeast paste and early L1 larvae were transferred to standard lab fly food with yeast paste after 24 hrs. After larvae enter wandering stage, the number of pupa was counted every 6 hours until all larvae pupariated.

### Pupal volume measurement

Pupae were picked from vials and imaged under dissection stereoscope. The length (L) and width (W) of pupae were measured using ImageJ software, and the pupal volume (V) was calculated in Microsoft Excel using the following equation,

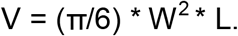

Volumes were then normalized to the average volume of control and the “Δ pupal volumes” were presented in figures.

### Fluorescence microscopy

Larvae were dissected in PBS and fixed using 3.7% formaldehyde for 15 mins at room temperature. Tissues were then washed in PBS and mounted in 90% glycerol for imaging. All confocal images were captured using Zeiss LSM710 confocal microscope.

### Immunohistochemistry

Larvae were dissected in PBS and fixed using 3.7% formaldehyde for 30 mins at room temperature. Tissues were washed in PBS containing 0.1% Triton X-100 (PBT) for 3 times and then permeabilized and blocked simultaneously using PBT containing 5% normal goat serum (NGS) for 1 hour. Tissues were then incubated with primary antibody (anti-Alk, 1:1000, anti-Pvr, 1:100, anti-phospho-Erk, 1:200) in PBT containing 10% NGS overnight at 4 degrees, followed by 5 washes and then post-secondary incubation for 2 hours at room temperature. DAPI staining occurred for 5 minutes at the pen-ultimate washing step after secondary antibody incubation. Finally, tissues were transferred to 70% glycerol/PBS mounting medium and then mounted on glass slide for imaging. Images were captured using a Zeiss LSM 710 confocal microscope.

### Ecdysteroid titer measurement

The ecdysteroid titers of larvae were measured using the 20-hydroxyecdysone Enzyme Immunoassay (EIA) kit (Cayman Chemicals), which detects both ecdysone (E) and 20-hydroxyecdysone (20E). Briefly, frozen larvae were homogenized in methanol and ecdysteroids were extracted as described previously (Warren et al., 2006). The extracts were evaporated in a Speed Vac and the residue resuspended in EIA buffer and analyzed following the manufacturer’s protocol. A standard curve was determined using a dilution series containing a known amount of purified 20E solution provided by the kit. Absorbance at 415 nm was detected using a benchtop microplate reader (Bio-Rad).

### Quantitative RT-PCR (qPCR)

Larvae were washed in PBS and then homogenized in Trizol (Invitrogen). Total RNA was purified using RNeasy Mini Kit (Qiagen) and cDNA library was obtained using SuperScript-III (Invitrogen) following the manufacturer’s protocol. qPCR was then carried out using SYBR Green reagent (Roche) on a LightCycler 480 platform. *Rpl23* was used as internal control for normalization. Primers used in this study are listed below.

*Rpl23* F 5’-GACAACACCGGAGCCAAGAACC -3’

R 5’-GTTTGCGCTGCCGAATAACCAC -3’

*nvd* F 5’-GGAAGCGTTGCTGACGACTGTG -3’

R 5’-TAAAGCCGTCCACTTCCTGCGA -3’

*spok* F 5’-TATCTCTTGGGCACACTCGCTG -3’

R 5’-GCCGAGCTAAATTTCTCCGCTT -3’

*sro* F 5’-CCACAACATCAAGTCGGAAGGAGC -3’

R 5’-ACCAGGCGAATGGAATCGGG -3’

*Cyp6t3* F 5’-GGTGTGTTTGGAGGCACTG -3’

R 5’-GGTGCACTCTCTGTTGACGA -3’

*phm* F 5’-GGATTTCTTTCGGCGCGATGTG -3’

R 5’-TGCCTCAGTATCGAAAAGCGGT -3’

*dib* F 5’-TGCCCTCAATCCCTATCTGGTC -3’

R 5’-ACAGGGTCTTCACACCCATCTC -3’

*sad* F 5’-CCGCATTCAGCAGTCAGTGG -3’

R 5’-ACCTGCCGTGTACAAGGAGAG -3’

### Quantification of autophagic vesicles

The number and area of Atg8a positive vesicles were quantified using imageJ software. Briefly, the vesicles were selected using the “threshold” function. Then the number and total area of the vesicles were calculated automatically using the “analyze particles” function in the software.

### Statistics

GraphPad Prism software was used to carry out statistical analyses. Student’s t-test was used to determine statistical significance.

