## Supplementary material for "Coordination Among Multiple Receptor Tyrosine Kinase Signals Controls *Drosophila* Developmental Timing and Body Size": Suppl figures and text

| **Gene name** | **Genome-wide screen** | **Targeted screen**  **(50% Pupariation)** | **P value** |
| --- | --- | --- | --- |
| Control |  | 4.794 ± 0.059 (Days AED) | N/A |
| **Alk** | NOP | **5.253 ± 0.055** | **0.0013** |
| btl | NOP | 4.727 ± 0.061 | 0.3281 |
| Cad96Ca | NOP | 4.704 ± 0.048 | 0.1703 |
| CG10702 | NOP | 4.714 ± 0.060 | 0.2483 |
| Ddr | NOP | 4.817 ± 0.101 | 0.7998 |
| dnt | NOP | 4.741 ± 0.046 | 0.3779 |
| drl | NOP | 4.673 ± 0.046 | 0.0874 |
| Drl-2 | NOP | 4.722 ± 0.033 | 0.2243 |
| **Egfr** | NOP | **L3 arrest** | N/A |
| Eph | NOP | 4.806 ± 0.032 | 0.8075 |
| htl | NOP | 4.488 ± 0.035 | **0.0061** |
| **InR** | **Major delay** | **> 5-day delay / L3 arrest** | N/A |
| Nrk | NOP | 4.528 ± 0.075 | **0.0186** |
| otk | NOP | 4.703 ± 0.080 | 0.2701 |
| **Pvr** | **Major delay** | 4.897 ± 0.068 | 0.1796 |
| Ret | NOP | 4.687 ± 0.055 | 0.1332 |
| Ror | NOP | 4.741 ± 0.050 | 0.387 |
| sev | NOP | 4.926 ± 0.048 | 0.0707 |
| Tie | NOP | 4.484 ± 0.080 | 0.0139 |
| **tor** | **Delay** | **6.302 ± 0.126** | **0.0008** |

**Table S1. Targeted screen for RTKs regulating developmental timing in the PG.**

The targeted screen was carried out using RNAi lines from TRiP (available from BDSC, for details of the RNAi lines used in the screen see the STAR Methods). Expression of the UAS-RNAi was driven by *phm-Gal4*. The 50% pupariation time of each group (Mean ± SEM, n=3) was compared with *phm>w1118* control and the p values (unpaired student’s t-test) were shown. A genome-wide RNAi screen for developmental timing regulators (Danielsen et al., 2016), which used RNAi lines from the VDRC, was also surveyed in this study and results of RTKs are listed in the table. “NOP” indicates no obvious phenotype on developmental timing; “delay” and “major delay” indicate 2-day and 3-day delay compared with control, respectively. The RNAi lines which cause developmental delay in at least one screen are marked in bold.

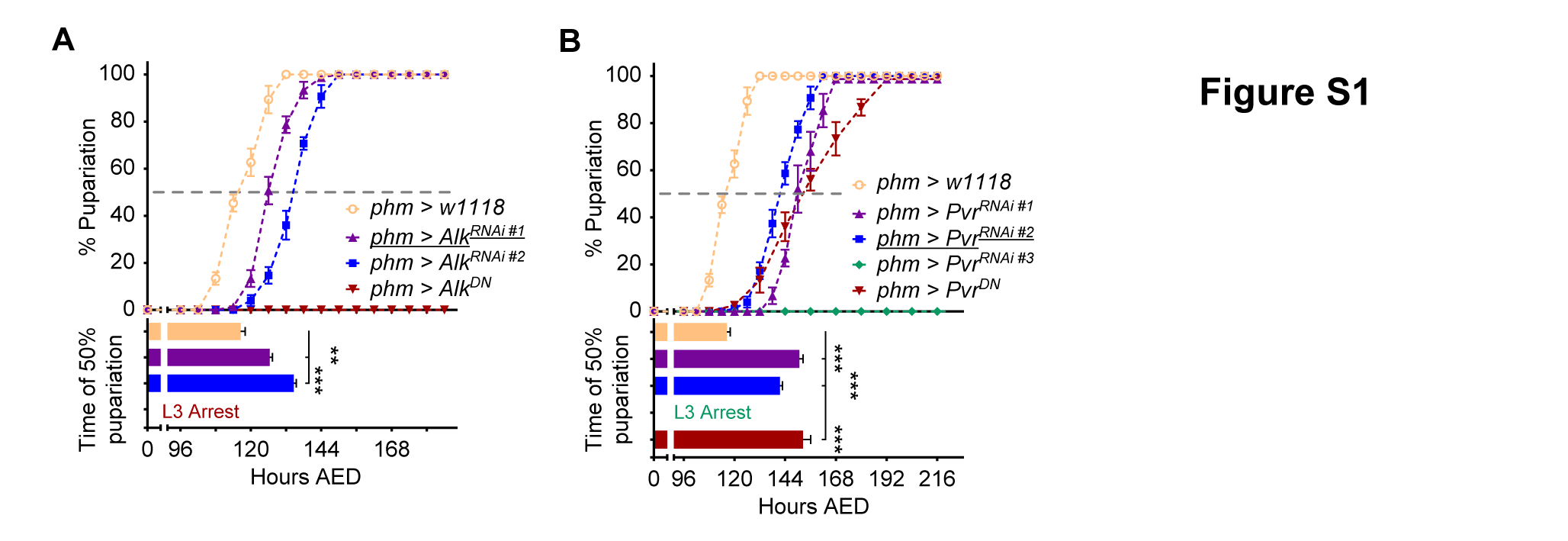

**Figure S1. Knockdown of Alk/Pvr in the PG causes delay/arrest of development. Related to Figure 1.**

(A and B) Pupariation timing curves and the time of 50% pupariation of Alk (A) and Pvr (B) knockdown larvae. Mean ± SEM; p values by unpaired t-test (n=3; **p<0.01, ***p<0.001). *UAS-Alk^RNAi^* lines tested are: #1, JF02668, TRiP; #2, v107083, VDRC. *UAS-Pvr^RNAi^* lines tested are: #1, v43459, VDRC; #2, v43461, VDRC; #3, v105353, VDRC. The underscored lines in the figure were used in the following study and were marked as *Alk^RNAi^* and *Pvr^RNAi^*, respectively.

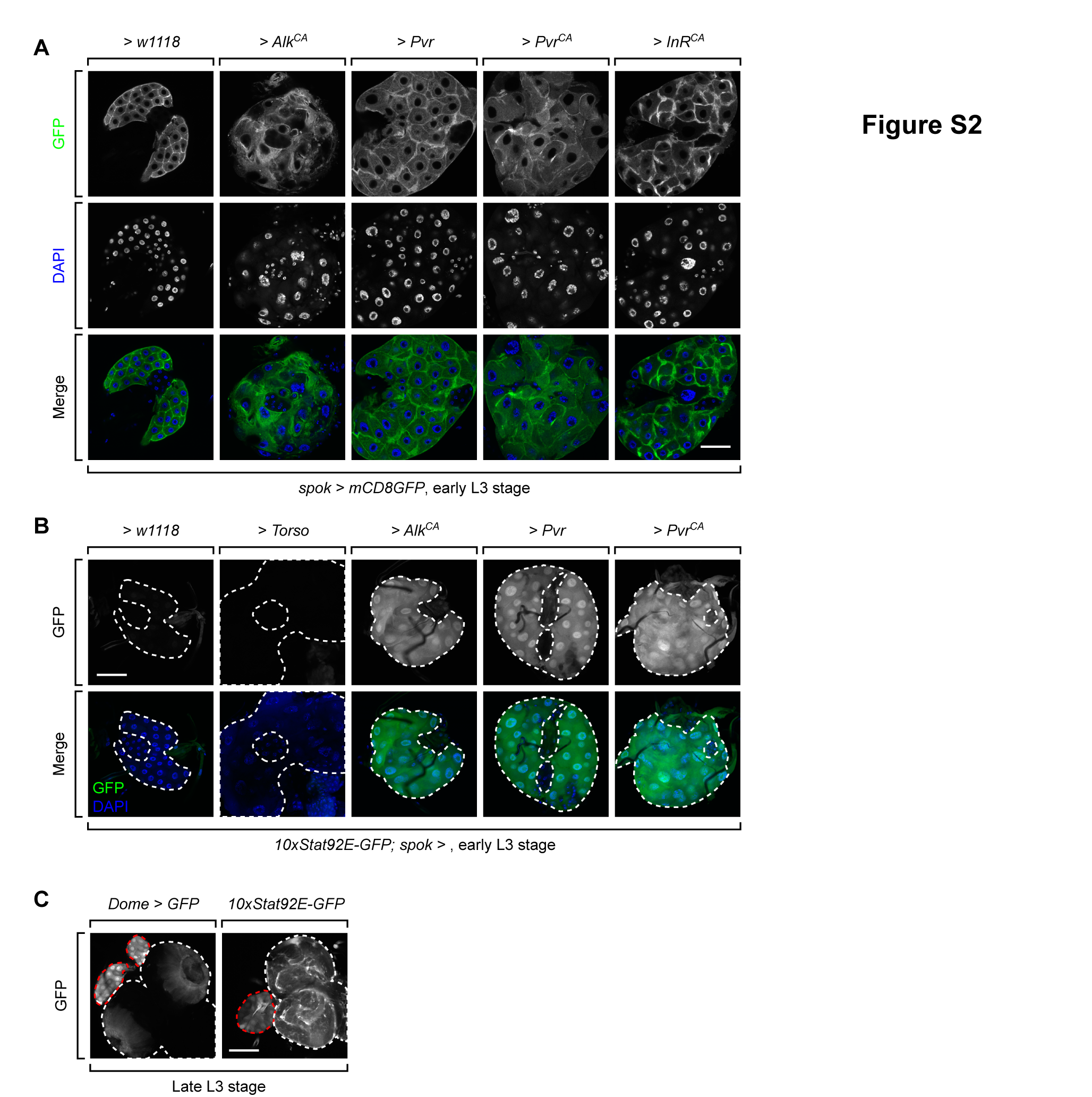

**Figure S2. Activation of Alk/Pvr causes overgrowth of PG with atypical morphology and activation of the Jak/Stat pathway. Related to Figure 4.**

(A) Images of *spok>w1118*, *spok>Alk^CA^*, *spok>Pvr*, *spok>Pvr^CA^* and *spok>InR^CA^* PGs expressing *mCD8GFP*. Cell membranes are marked by mCD8GFP and atypical morphology can be observed in *spok>Alk^CA^* and *spok>Pvr^CA^* PGs. Scale bar, 50μm. (B) Images of *spok>w1118*, *spok>Torso*, *spok>Alk^CA^*, *spok>Pvr* and *spok>Pvr^CA^* PGs with *10xStat92E-GFP* reporter. Dash lines outline the PG region in the ring glands. Scale bar, 50μm. (C) Images of ring gland-brain complexes of wildtype late-L3 larvae. GFP expression is controlled by either *Dome-Gal4* (left) or *10xStat92E-GFP* reporter (right). White dash lines mark the brain lobe and ventral ganglia, while red dash lines mark the PG region. Scale bar, 100μm.

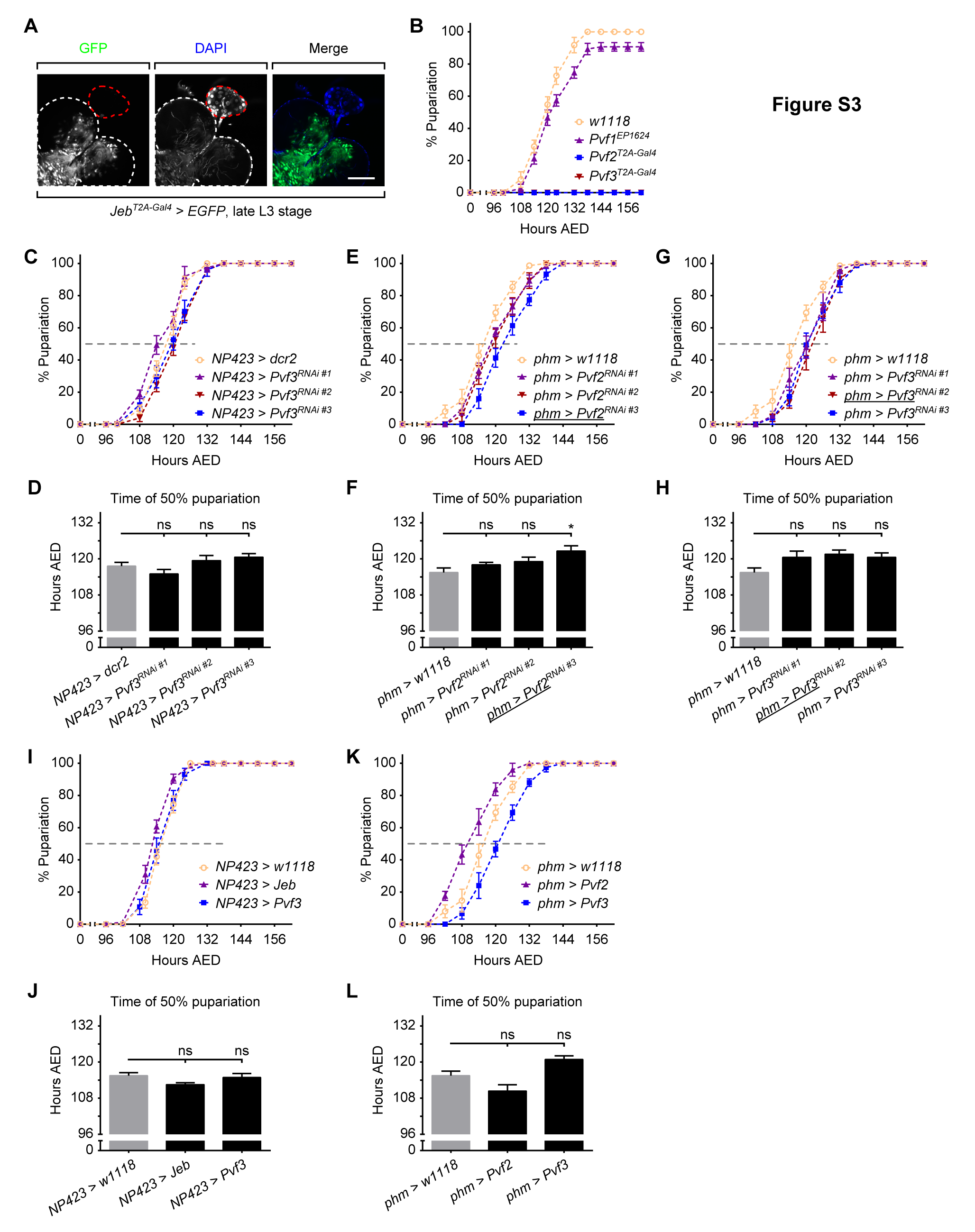

**Figure S3. Ligands that activate Alk and Pvr derive from both PGNs and PG. Related to Figure 5.**

(A) Images of the brain-ring gland complex of *Jeb^T2A^>EGFP* larvae. White dash lines mark the brain lobe and ventral ganglia, while red dash lines mark the PG region. Scale bar, 100μm. (B) Pupariation timing curves of *Pvf1*, *Pvf2* and *Pvf3* mutants. A well characterized *Pvf1^EP1624^* was used as *Pvf1* null mutant, while the T2A Gal4 lines were tested as *Pvf2* and *Pvf3* null mutants. (C and D) Pupariation timing curves (C) and the time of 50% pupariation (D) of larvae with Pvf3 knockdown in the PGNs. (E-H) Pupariation timing curves (E and G) and the time of 50% pupariation (F and H) of larvae with Pvf2 (E and F) and Pvf3 (G and H) knockdown in the PG. *UAS-Pvf2^RNAi^* lines tested are: #1, HMJ23540, TRiP; #2, 13780R-2, NIG; #3, v7629, VDRC. *UAS-Pvf3^RNAi^* lines tested are: #1, HMS01876, TRiP; #2, 13781R-1, NIG; #3, v37933, VDRC. (I and J) Pupariation timing curves (I) and the time of 50% pupariation (J) of larvae with Jeb and Pvf3 overexpression in the PGNs. (K and L) Pupariation timing curves (K) and the time of 50% pupariation (L) of larvae with Pvf2 and Pvf3 overexpression in the PG. (D, F, H, J and L) Mean ± SEM; p values by unpaired t-test (n=3; ns, not significant, *p<0.05).
